# Xenopus IgX informs engineering strategies of IgM and IgG hexamers

**DOI:** 10.1101/2025.07.23.666482

**Authors:** Ruixue Zhang, Chenggong Ji, Shuhan Li, Ningning Li, Ning Gao, Junyu Xiao

**Author notes:** Corresponding author. (C.J.); (N.G.); (J.X.). These authors contributed equally to this work (Ruixue Zhang, Chenggong Ji).

## Abstract

Polymeric immunoglobulins are essential components of the immune system in jawed vertebrates. While mammalian immunoglobulin M (IgM) typically forms a pentamer linked by the joining chain (J-chain), *Xenopus laevis* IgX assembles into a J-chain-independent polymer. Here, we present the cryo-electron microscopy (cryo-EM) structure of IgX, revealing its hexameric configuration. By incorporating the IgX tailpiece into human IgM, we achieved efficient IgM hexamer formation. Truncating IgM’s natural tailpiece to a range of 11-16 residues also substantially enhanced hexamerization efficiency. Furthermore, introducing a shortened IgM tailpiece to IgG resulted in effective IgG hexamer formation. We further show that the engineered IgM and IgG hexamers targeting CD20 demonstrated robust complement-dependent cytotoxicity (CDC) against several B lymphoma cells. Additionally, the IgG-Fc hexamer functioned as a decoy, attenuating CDC in cell cultures. These findings deepen our understanding of polymeric immunoglobulin evolution and introduce innovative strategies for the development of IgM- and IgG-based biologics.

**One Sentence Summary:** The cryo-EM structure of the African clawed frog immunoglobulin IgX reveals a uniform hexameric assembly, inspiring innovative strategies for the development of therapeutic human IgM and IgG hexamers.

## Introduction

Polymeric immunoglobulins (pIgs) play an essential role in the immune systems of all jawed vertebrates(*1, 2*). Immunoglobulin M (IgM) is one of the most ancient classes of antibodies, preserved throughout evolutionary history and serving as a prototype for pIgs. Mammalian IgM typically assembles into a pentamer when combined with the joining chain (J-chain)(*3–5*). The J-chain imparts unique features to the IgM pentamer and enhances its interactions with various receptors and binding partners such as pIgR, FcμR, and CD5L(*3, 6–8*). An 18-residue tailpiece at the C-terminus of the IgM heavy chain is crucial for polymerization and association with the J-chain. A similar tailpiece is found in IgA, promoting its polymerization and interaction with the J-chain as well(*9–11*). Without the J-chain, mammalian IgMs can form multiple polymeric structures, including hexamers, pentamers, and tetramers(*12–18*). Interestingly, J-chain is absent in teleost fish, and teleost IgM exhibits a distinctive tetrameric configuration(*19–21*).

The multivalent nature of pIgs confers high avidity, crucial for both binding antigens with high affinity and triggering specific antibody effector functions. IgM, for instance, is a potent activator of the complement system, an essential component of the innate immune system that facilitates the elimination of microbial pathogens and damaged cells(*22, 23*). The classical complement pathway is initiated by the activation of the C1 complex, with C1q binding to IgM or IgG complexed with antigens. Notably, the IgM hexamer is significantly more effective in activating the complement system than the pentamer, likely due to the hexameric structure of the C1q adaptor(*15, 17, 24–26*). IgG also activates the classical complement pathway; however, since IgG is a monomer in solution, clustering of multiple IgG molecules, ideally six, is necessary for efficient C1q recruitment(*27–30*).

Engineering IgG to promote hexamer formation has become an important focus in the development of antibody therapeutics, given the high complement-dependent cytotoxicity (CDC) activities exhibited by these engineered molecules(*31*). Strategies to create IgM-like IgG have included fusing the 18-amino acid IgM tailpiece (μtp)(*32, 33*), or the C575S variant(*34*), to the heavy chain of IgG, exploiting the polymerization capabilities of the μtp. Additionally, the L309C mutation has been introduced into IgG-Fc (Fcγ) to mimic Cys414 in IgM, promoting disulfide bond formation between adjacent IgG molecules(*32, 33, 35*). Further modifications include altering the C-terminal six amino acids of Fcγ (SLSPGK) to the IgM sequence preceding μtp (DKSTGK) and introducing two point mutations, V567I and A572G, into the μtp to improve polymerization (denoted as Fcγ_Innovent_ hereafter)(*36*). Moreover, IgG hexamer structures have been identified in the crystal lattices of several antibodies, including two anti-HIV-1 gp120 antibodies, b12 and 2G12(*37, 38*). Drawing on the reconstructed hexamer structure of b12 from crystal symmetry, point mutations such as E345R, E430G, and S440Y have been strategically designed to enhance Fcγ-Fcγ interactions and promote hexamer formation(*28, 39*). These innovations underpin the HexaBody antibody technology platform, marking a pivotal advancement in antibody engineering. Similarly, mutations of H429Y or H429F (Stellabody)(*40*), as well as Q311R/M428E/N434W(*41*), also result in increased IgG oligomer formation and/or complement activation.

The African clawed frog, *Xenopus laevis*, serves as an evolutionary link between teleost fish and more complex vertebrates and is an ideal model for studying early diversification events in immunoglobulins. In Xenopus, two types of pIgs, IgM and IgX, are present. IgX is predominantly found in mucus secretions and is thought to be a functional analogue of IgA(*42, 43*); although it shares more structural similarities with IgM, having four constant domains compared to the three in IgA(*44*). IgX forms a polymer(*45*); but unlike IgM and IgA, IgX does not appear to associate with the J-chain(*42*). This observation has prompted us to further investigate into its assembly mechanism.

## Results

### Xenopus IgX forms a hexamer

The Fc region of the IgX heavy chain (Fcχ) includes three constant domains and an 11-residue tailpiece (Cχ2, Cχ3, Cχ4, and χtp). We expressed Fcχ recombinantly using HEK293F cells. The polymeric form was subsequently separated by size-exclusion chromatography (SEC) (fig. S1). Cryo-EM 2D classifications revealed that the Fcχ polymer consistently forms hexamers (Fig. 1A). The fragment consisting of Cχ3, Cχ4, and χtp of Fcχ (Fcχ^Cχ3-Cχ4-χtp^) was also purified and found to be able to form a stable hexamer. This fragment displayed improved qualities on EM grids and was subjected to detailed cryo-EM analysis, achieving a 3D reconstruction at 3.3 Å resolution (Fig. 1B, fig. S2, and Table S1).

**Fig. 1.**
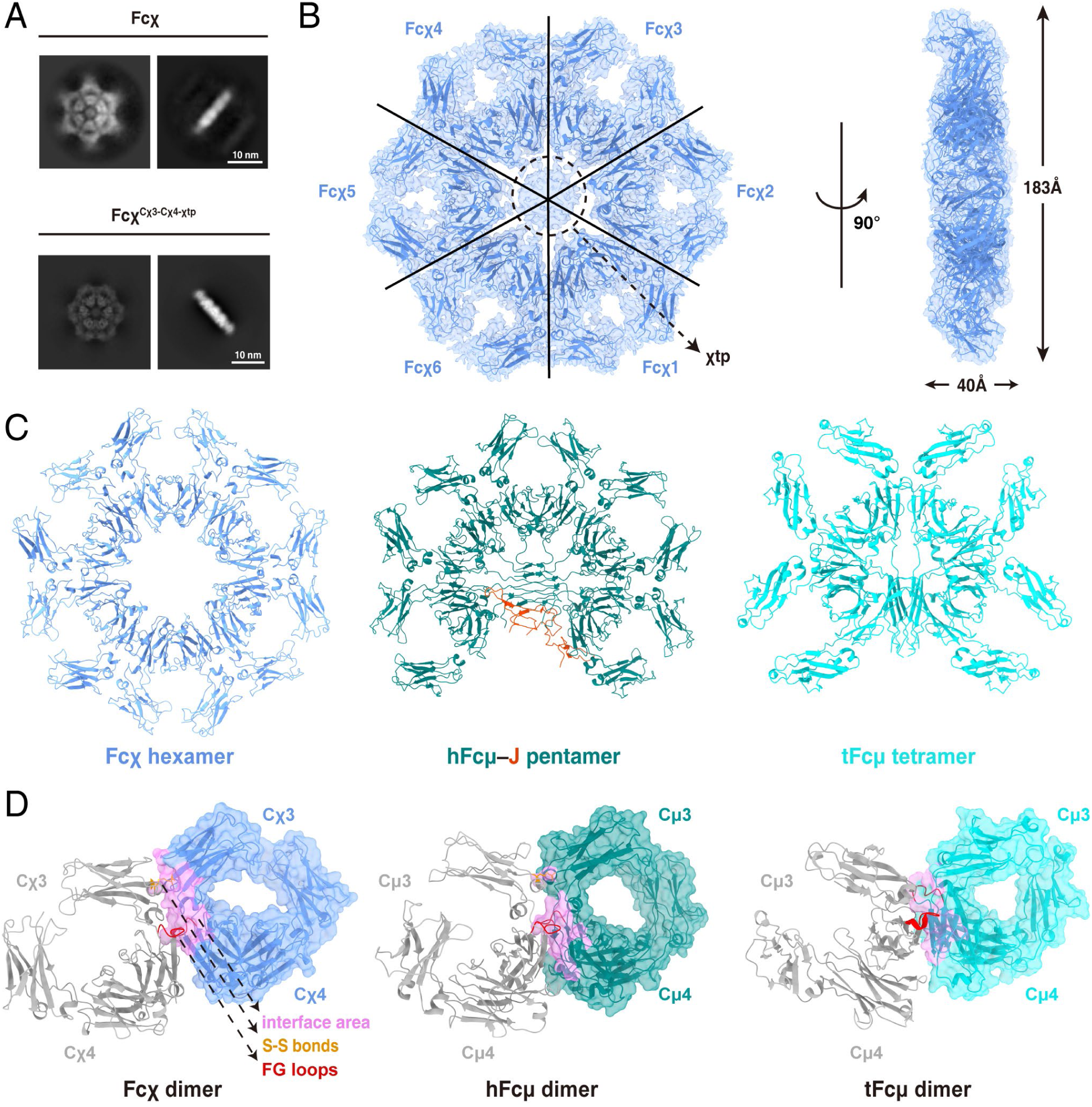
Xenopus Fcχ can form a stable hexamer. (A) Cryo-EM 2D analyses showed that both Fcχ and Fcχ^Cχ3–Cχ4–χtp^ form hexamers. (B) Cryo-EM reconstruction of the Fcχ^Cχ3–Cχ4–χtp^ hexamer at 3.29 Å resolution, shown in two orientations. The EM density map is superimposed on the structural model. The χtp regions were not clearly resolved, due to the blurred densities in this region, as indicated by a dashed circle. (C) Side-by-side comparisons of the Fcχ hexamer (blue), human Fcμ-J pentamer (Fcμ in teal and J-chain in orange), and teleost Fcμ tetramer (cyan). (D) Side-by-side comparisons of the Fc–Fc interactions in the Fcχ hexamer, human Fcμ-J pentamer, and teleost Fcμ tetramer. The interface area, the interchain disulfide bonds, and the FG loops of Cμ4 or Cχ4 domains are highlighted in pink, yellow, and red, respectively.

The Fcχ^Cχ3-Cχ4-χtp^ structure features six Cχ3-Cχ4 units aligned in the same plane, arranged in a hexagonal symmetry (Fig. 1C). In contrast, the human IgM pentameric core structure (Fcμ-J) displays five Fcμ molecules arranged in a pseudo-hexagonal symmetry pattern, with the J-chain filling the gap between the first and fifth Fcμ (Fig. 1C, fig. S3A). On the other hand, the teleost IgM-Fc (tFcμ) presents a different configuration, forming a tetramer with a more relaxed arrangement and lacking the hexagonal symmetry observed in the Fcχ^Cχ3-Cχ4-χtp^ and Fcμ-J structures (Fig. 1C, fig. S3B).

Further local refinements on the interface between adjacent Fcχ units enhanced the visualization of interface residues (Fig. 1D). Similar to the Fcμ-Fcμ interface in the human IgM structure, both Cχ3 and Cχ4 domains are involved in the Fcχ-Fcχ interactions. The Cχ4-Cχ4 interface between adjacent Fcχ units buries approximately 400 Å² of surface area from each Cχ4, comparable to the interface between two Cμ4 domains in Fcμ-J. Cys430 in the Cχ3 domain participates in forming interchain disulfide bonds, similar to Cys414 in the Cμ3 domain of Fcμ. In contrast, a similar Cys is absent in the teleost IgM (fig. S4), and the interface between adjacent tFcμ units in the tetramer is almost entirely contributed by the Cμ4 domains (Fig. 1D).

The χtp regions were poorly resolved in the EM density map refined with C1 symmetry (Fig. 1B). Given that the distinct hexagonal symmetry of the Cχ3-Cχ4 region likely played a dominant role in particle alignment compared to the asymmetric χtp region, the Fcχ^Cχ3–Cχ4–χtp^ particles were processed using C6 symmetry expansion (fig. S2A). Secondary structure features of the χtps became discernible in one of the resulting C6 symmetry-expanded classes (class 6). Although accurate model building was not feasible due to limited resolution, it is evident that the χtp region displays characteristic features of β-sheet conformation. The χtp shows strong sequence conservation with the tailpiece β-strand regions in human Fcμ and tFcμ (Fig. 2A), which form β-sandwich structures in both human IgM pentamers and teleost IgM tetramers, despite their distinct structural configurations (Fig. 1C). The removal of χtp severely impaired Fcχ hexamer formation, underscoring its essential role in the hexamerization of IgX (fig. S1A).

**Fig. 2.**
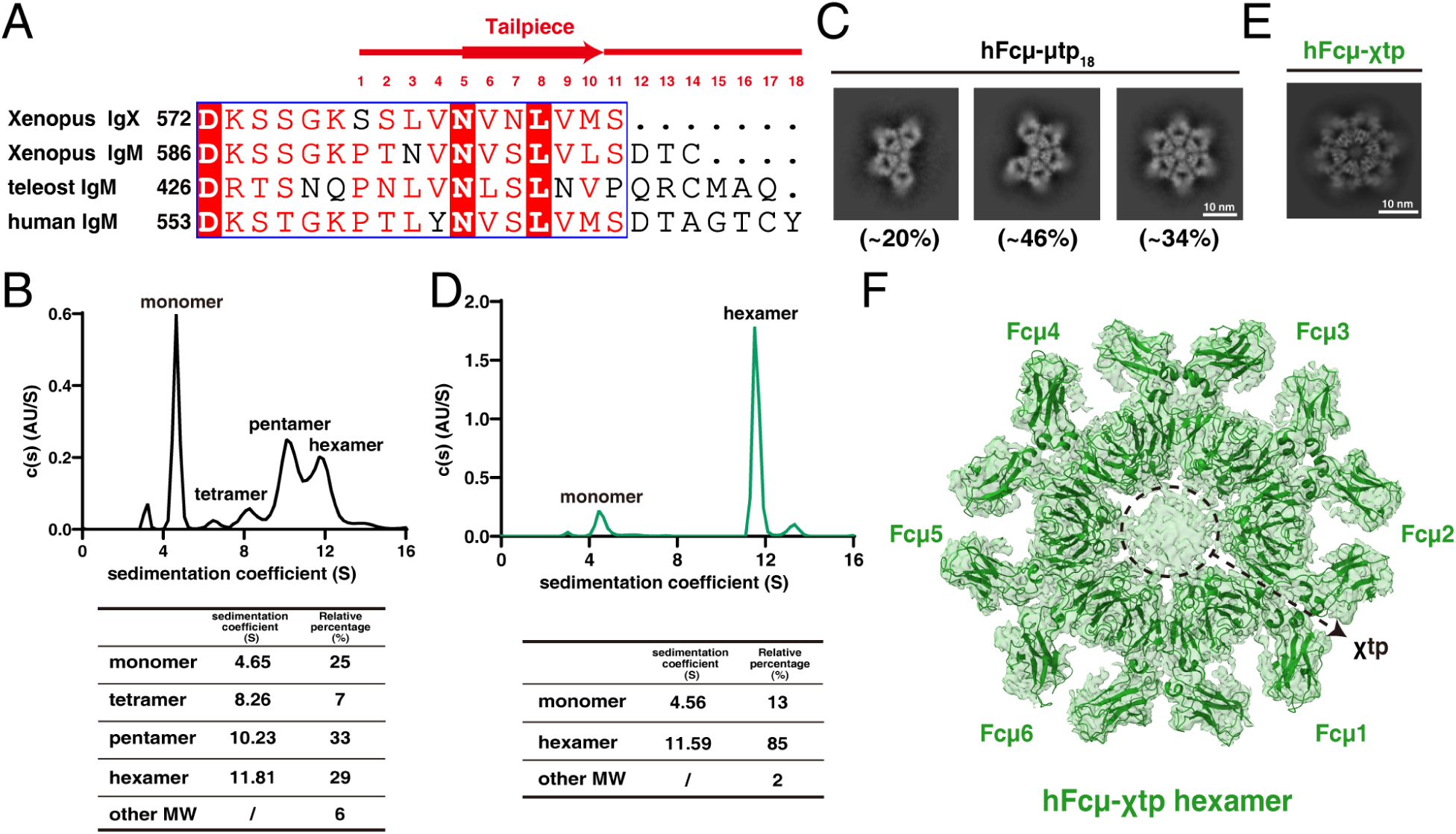
The Fcμ-χtp chimera forms a hexamer. (A) Sequence alignment of the tailpiece regions of Xenopus IgX and IgM, teleost IgM (from *Oncorhynchus mykiss*), and human IgM. The tailpiece regions are highlighted. (B) SV-AUC analysis of Fcμ-μtp_18_ suggested that monomer, tetramer, pentamer, and hexamer are all present in solution. (C) Cryo-EM 2D analyses suggested that the polymers of Fcμ-μtp_18_ consist of a mixture of tetramer, pentamer, and hexamer. (D) SV-AUC analysis indicated that the Fcμ–χtp polymer is uniformly hexameric, in contrast to Fcμ-μtp_18_. (E) Cryo-EM 2D analyses of the Fcμ-χtp chimera. (F) Cryo-EM reconstruction of the Fcμ-χtp hexamer at 3.29 Å resolution. The EM density map is superimposed on the structural model.

### The Fcμ-χtp chimera forms a uniform hexamer

Mammalian IgM molecules form stable pentamers in the presence of the J-chain; in its absence, however, IgM exhibits reduced stability and can assume various polymeric forms including hexamers, pentamers, and even tetramers(12–18). Indeed, recombinant Fcμ with an intact 18-amino acid tailpiece (denoted as Fcμ-μtp_18_) displays heterogeneity in solution, forming tetramers, pentamers, and hexamers as determined by sedimentation velocity analytical ultracentrifugation (SV-AUC) (Fig. 2B). These polymeric forms were enriched by SEC (fig. S1B) and analyzed using cryo-EM. Consistent with the SV-AUC results, the tetramer, pentamer, and hexamer forms of Fcμ-μtp_18_ were all distinctly identifiable in the 2D classes (Fig. 2C).

To achieve a uniform Fcμ hexamer, we took inspiration from the Fcχ hexamer structure. We first replaced the 18-amino acid tailpiece segment of Fcμ with the 11-amino acid χtp, creating a Fcμ-χtp chimera. This chimera was recombinantly expressed and purified by SEC (fig. S1, B and C). Interestingly, SV-AUC analysis suggested that the polymeric form of this chimera consistently displayed a uniform hexameric structure (Fig. 2D). The Fcμ-χtp polymer was further subjected to cryo-EM analysis, confirming the exclusive presence of a uniform hexameric assembly with no other polymeric forms detected (Fig. 2E). The cryo-EM structure of the Fcμ-χtp hexamer was subsequently resolved at a resolution of 3.3 Å (Fig. 2F, fig. S5, and Table S1). Similar to the Fcχ hexamer, the six Fcμ units in this structure are arranged in a perfect hexagonal geometry (fig. S3C).

### Shortened tailpieces facilitate IgM hexamer formation

Compared to the 18-amino acid tailpiece of Fcμ (μtp_18_), the χtp comprises only 11 amino acids (Fig. 2A). This difference led us to hypothesize that the length of the tailpiece influences the assembly of Fcμ hexamers. To test this hypothesis, we first created the Fcμ-μtp_11_ variant by truncating the last seven amino acids from the C-terminus of μtp. This variant was recombinantly purified using the same methods as those for Fcχ and the Fcμ-χtp chimera (fig. S1, B and C). SV-AUC analysis revealed that the Fcμ-μtp_11_ polymers are homogeneous hexamers (Fig. 3A), which is confirmed by cryo-EM imaging (Fig. 3B).

**Fig. 3.**
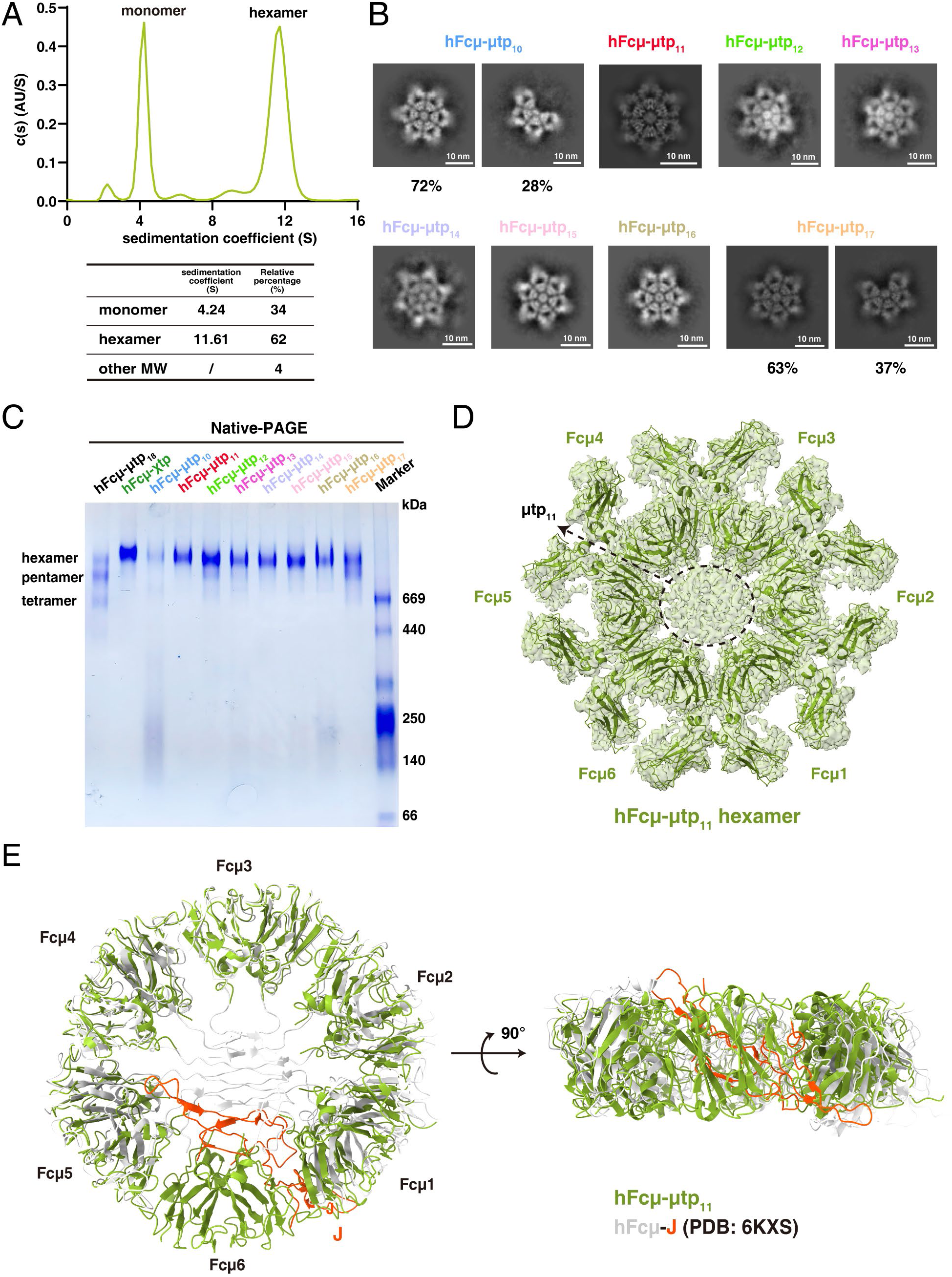
Human Fcμ variants with shorter tailpieces can form hexamers. (A) SV-AUC analysis suggested besides monomer, Fcμ-μtp_11_ is only present as a hexamer in solution. (B) Cryo-EM 2D analyses of Fcμ-μtp variants with different tailpiece lengths. (C) Native-PAGE analyses of the Fcμ variants. (D) Cryo-EM reconstruction of the Fcμ-μtp_11_ hexamer at 3.29 Å resolution. The EM density map is superimposed on the structural model. (E) A structural comparison between the Fcμ-μtp_11_ hexamer and Fcμ-J pentamer focusing on the Cμ4 regions. The Fcμ molecules in the Fcμ-μtp_11_ hexamer are represented in green, while those in the Fcμ-J pentamer are shown in white. The J-chain is highlighted in orange.

To further investigate the impact of tailpiece length on the hexamerization of Fcμ, we systematically truncated the μtp by sequentially removing amino acids from the C-terminus (fig. S1C). The multimer fractions of each variant, collected after SEC, were analyzed using cryo-EM 2D classifications (Fig. 3B). The analysis revealed that μtp lengths ranging from 11 to 16 amino acids consistently resulted in the formation of Fcμ hexamers. Notably, the Fcμ-μtp_10_ variant exhibited both hexamers and tetramers, while Fcμ-μtp_17_ displayed a mix of hexamers and pentamers. The oligomeric states of these multimers were further confirmed by native polyacrylamide gel electrophoresis (Native-PAGE) analyses: Fcμ-μtp_10_, Fcμ-μtp_17_, and Fcμ-μtp_18_ formed heterogeneous polymers in solution (Fig. 3C). In contrast, Fcμ-χtp, as well as Fcμ-μtp_11_ to Fcμ-μtp_16_, appeared as homogeneous hexamers. Together, these results suggest that a tailpiece containing 11-16 residues optimally supports the hexamer formation of Fcμ.

The cryo-EM structure of the Fcμ-μtp_11_ hexamer was further resolved at an overall resolution of 3.2 Å, enabling clear visualization of all six Fcμ molecules, particularly the Cμ4 domains (Fig. 3D, fig. S6). All six Fcμ units are arranged in the same plane, with the 6th Fcμ molecule integrated into the hexamer by occupying the position of the J-chain in the Fcμ-J pentamer (Fig. 3E), consistent with previous predictions. When the Fcμ-J pentamer core, including five pairs of Cμ4 and the J-chain, is superimposed onto the Fcμ hexamer, Fcμ1 and Fcμ5 in the pentamer exhibit slight rotations due to their direct associations with the J-chain (Fig. 3E, fig. S3D). This delicate asymmetric feature, introduced by the J-chain, impacts interactions between the IgM pentamer and its specific receptor FcμR(6).

### Engineered IgG with μtp_11_ assembles into a hexamer

To further investigate whether μtp_11_ can also lead to IgG hexamer formation, we introduced the μtp_11_ segment into the C-terminus of human IgG1-Fc (Fcγ), creating the Fcγ-μtp_11_ chimera (Fig. 4A). Additionally, we replaced the C-terminal six amino acids of Fcγ (SLSPGK) with the linker sequence between Cμ4 and μtp (DKSTGK), inspired by the Fcγ_Innovent_ design(36). This replacement removes the rigid Pro445 in Fcγ, potentially increasing the flexibility in the linker region between Fcγ and μtp_11_.

**Fig. 4.**
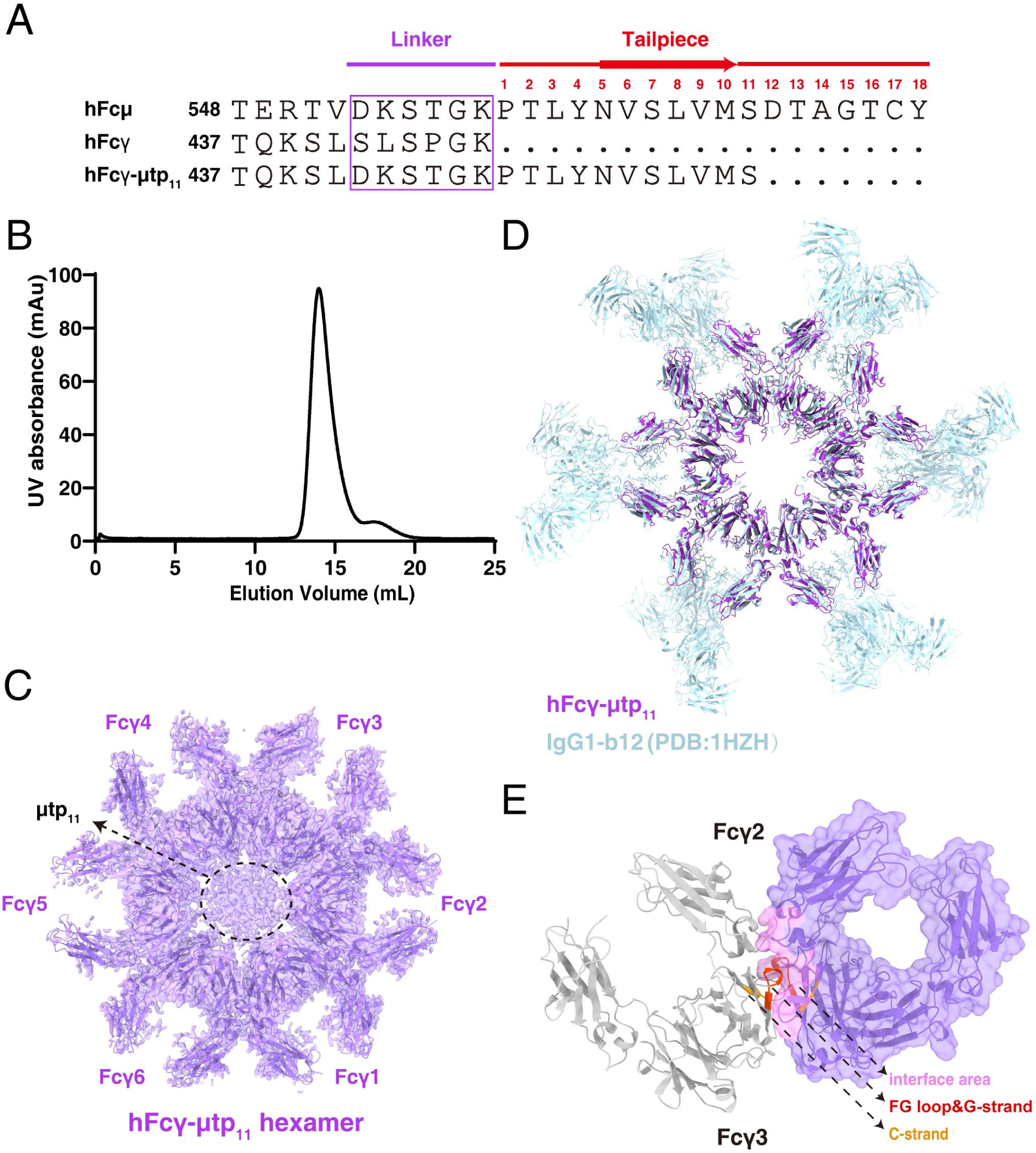
The hFcγ–μtp_11_ hexamer. (A) Sequences of Fcμ, Fcγ, and the engineered Fcγ-μtp_11_ chimera at the C-terminal regions. (B) SEC analysis suggested that Fcγ–μtp_11_ is mainly present as a hexamer in solution. (C) Cryo-EM reconstruction of the Fcγ-μtp_11_ hexamer at 2.75 Å resolution. (D) Structural overlay of the Fcγ-μtp_11_ hexamer and the IgG1-b12 hexamer within the crystal lattice (PDB ID: 1HZH). The Fcγ-μtp_11_ hexamer is shown in purple, while the IgG1-b12 hexamer is depicted in light cyan. (E) Fc–Fc interactions in the Fcγ hexamer. The interface area in one of Fcγ protomer is highlighted in pink. Important regions in the Cγ3 domains that are involved in the interactions between two adjacent Fcγ molecules, including the C-strand, and the FG loop & G-strand, are highlighted in red and yellow.

The assembly of the Fcγ-μtp_11_ polymer was highly efficient, as indicated by the SEC analysis (Fig. 4B). The cryo-EM structure of the Fcγ-μtp_11_ hexamer was subsequently determined at a resolution of 2.75 Å (Fig. 4C, fig. S7). Six Fcγ molecules cluster into a hexagon in the structure, displaying high similarity to the b12 hexamer reconstructed from crystal symmetry (Fig. 4D). Both the Cγ2 and Cγ3 domains contribute to the interaction between adjacent Fcγ molecules, with approximately 960 Å² of surface area from each molecule buried in the interface (Fig. 4E). Notably, residues Glu380 and Glu382 in the C-strand, along with the His433-Gln438 segment in the FG loop and G-strand of the Cγ3 domain, play pivotal roles in mediating the Fcγ-Fcγ interactions (fig. S7, E and F).

### Enhanced CDC activities of the IgM and IgG hexamers with μtp_11_

To evaluate the potential of engineered hIgM and hIgG hexamers to enhance complement activation, we conducted complement-dependent cytotoxicity (CDC) experiments. The heavy chain region of the antigen-binding fragment of rituximab (RTX, denoted as RTX-Fcγ hereafter), an anti-CD20 IgG, was engineered into Fcμ, Fcμ-μtp_11_, and Fcγ-μtp_11_ constructs. We then co-expressed these constructs with the light chain of RTX and, for the IgM pentamer, the J-chain as well, resulting in recombinant anti-CD20 IgM pentamer (RTX-Fcμ-J), IgM hexamer (RTX-Fcμ-μtp_11_), and IgG hexamer (RTX-Fcγ-μtp_11_) (fig. S1D). For comparison, we also prepared recombinant RTX-Fcγ_E430G_ and RTX-Fcγ_Innovent_.

CDC activity was first assessed using the OCI-Ly10 B lymphoma cells in the presence of human complement(46). As anticipated, the monomeric RTX-Fcγ showed the lowest complement-activating activity among these molecules, with an IC50 of approximately 0.50 μg/mL (Fig. 5A). In contrast, the engineered RTX-Fcμ-J pentamer demonstrated an IC50 of about 0.23 μg/mL, indicating a two-fold increase in CDC activity. The RTX-Fcμ-μtp_11_ hexamer showed a further two-fold increase in CDC activity compared to the RTX-Fcμ-J pentamer, with an IC50 of 0.12 μg/mL. Importantly, both RTX-Fcγ_E430G_ and RTX-Fcγ_Innovent_ exhibited strong complement-activating capabilities, with IC50 values of approximately 0.06 μg/mL, nine times more potent than the original RTX-Fcγ (Fig. 5A); and the CDC activity of our RTX-Fcγ-μtp_11_ design was comparable to that of RTX-Fcγ_Innovent_ and RTX-Fcγ_E430G_.

**Fig. 5.**
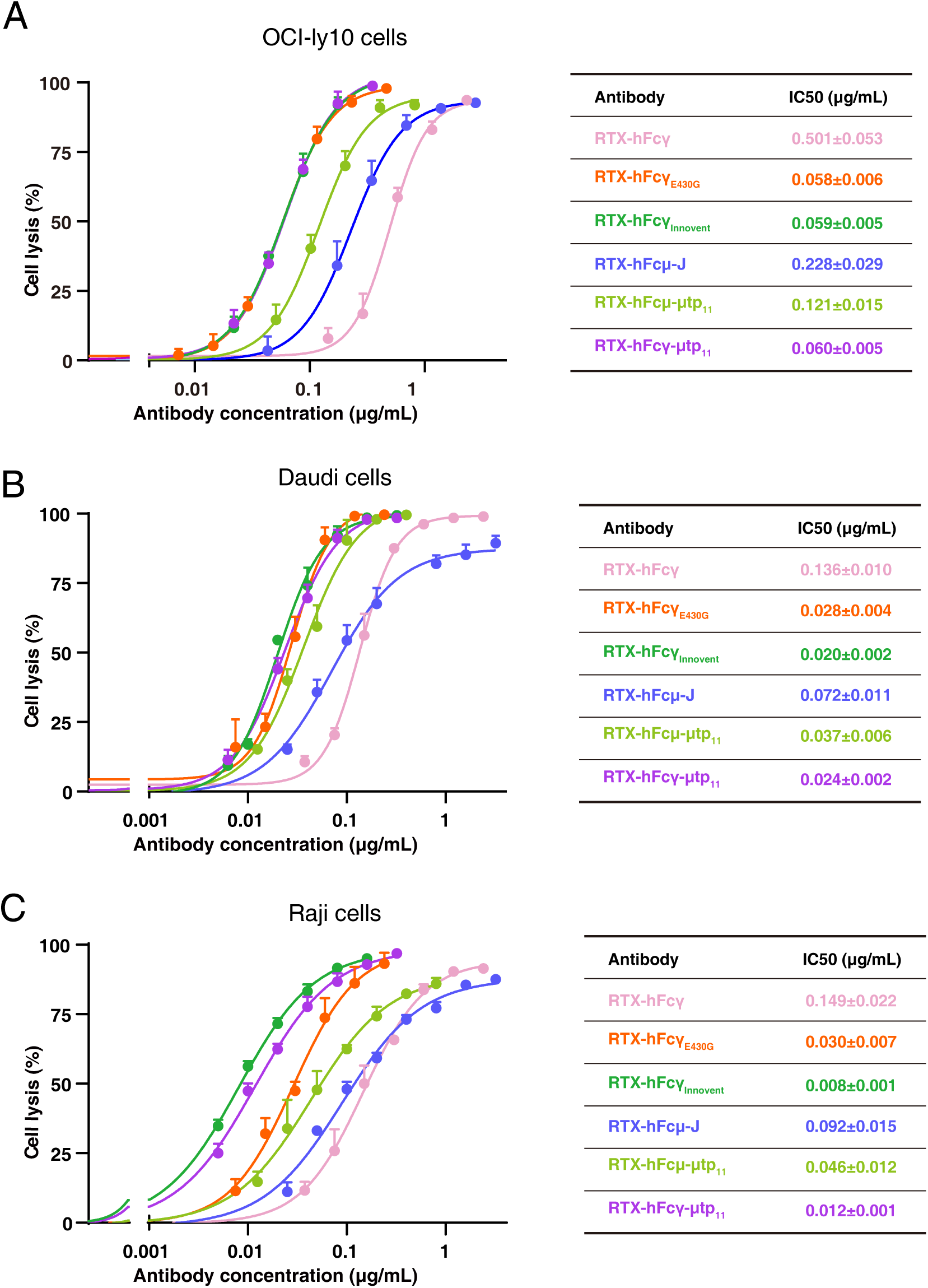
IgM and IgG hexamers with μtp_11_ display enhanced CDC activities. (A) CDC activities of the engineered RTX-IgG and RTX-IgM antibodies were assessed using OCI-Ly10 cells in the presence of human complement. Data were plotted as the mean ± 1 s.d. *n* = 3 biological replicates. Source data are provided in the Source Data file. (B) CDC activities of the engineered RTX-IgG and RTX-IgM antibodies towards the Daudi cells. To facilitate the assay, the Daudi cells, and also the Raji cells in c, were first cultured for several days in the RPMI-1640 medium supplemented with 10% heat-inactivated FBS. Data were plotted as the mean ± 1 s.d. *n* = 3 biological replicates. (C) CDC activities of the engineered RTX-IgG and RTX-IgM antibodies towards the Raji cells. Data were plotted as the mean ± 1 s.d. *n* = 3 biological replicates.

We also measured the CDC activity of these engineered molecules in other B lymphoma cell lines, including Daudi and Raji (Fig. 5, B and C). Unlike the OCI-Ly10 cells, which were cultured with standard fetal bovine serum (FBS) and used directly in the CDC assay, the Daudi and Raji cells were first cultured with heat-inactivated FBS prior to the assay. This step appeared to enhance their sensitivity to CDC-mediated killing. Nevertheless, the engineered molecules exhibited a generally consistent pattern of activity in Daudi cells (Fig. 5B), with potencies in the following order: RTX-Fcγ monomer < RTX-Fcμ-J pentamer < RTX-Fcμ-μtp_11_ hexamer < RTX-Fcγ_E430G_ ≈ RTX-Fcγ-μtp_11_ ≈ RTX-Fcγ_Innovent_. Notably, the RTX-Fcμ-J pentamer exhibited a shallower concentration-dependent response compared to the RTX-Fcμ-μtp_11_ hexamer in these cells. The differential potency of hexameric and pentameric IgM for complement activation is influenced by antigen densities(26), which may account for the different patterns observed between the RTX-Fcμ-J pentamer and the RTX-Fcμ-μtp_11_ hexamer in OCI-Ly10 and Daudi cells. In Raji cells, these molecules exhibited comparable activities to those observed in Daudi cells, with a notable difference that the RTX-Fcγ_Innovent_ and RTX-Fcγ-μtp_11_ hexamers exerted more potent CDC activities than RTX-Fcγ_E430G_ (Fig. 5C).

Collectively, these results suggest that the IgM and IgG hexamers engineered using the μtp_11_ strategy display substantially enhanced complement activation capabilities.

### Fcγ-μtp_11_ blocks complement activation

Excessive complement activation is associated with a range of diseases, and a recombinant Fc hexamer may serve as a “decoy” therapeutic to mitigate complement-mediated tissue damage. For example, a recent study showed that CSL777, a recombinant Fcγ-μtp_18_ fusion protein with the L309C mutation(35), provides protective effects in a mouse model of alloantibody-mediated acute lung injury(47). To investigate the effects of Fcγ-μtp_11_ and Fcμ-μtp_11_ hexamers on complement activation, we assessed the complement-dependent cytotoxicity (CDC) activity of RTX-Fcγ in Daudi and Raji cells in the presence of these proteins. Our results indicate that the Fcγ-μtp_11_ hexamer effectively inhibits complement activation mediated by RTX-Fcγ in both cell lines (Fig. 6, A and B). In contrast, the Fcμ-μtp_11_ hexamer showed no inhibitory effects. These findings are consistent with previous structural analyses, which demonstrated that the engineered IgG hexamer readily interacts with the complement C1 complex in solution, while substantial conformational changes upon antigen binding are required in IgM to expose the C1 binding sites(25, 29).

**Fig. 6.**
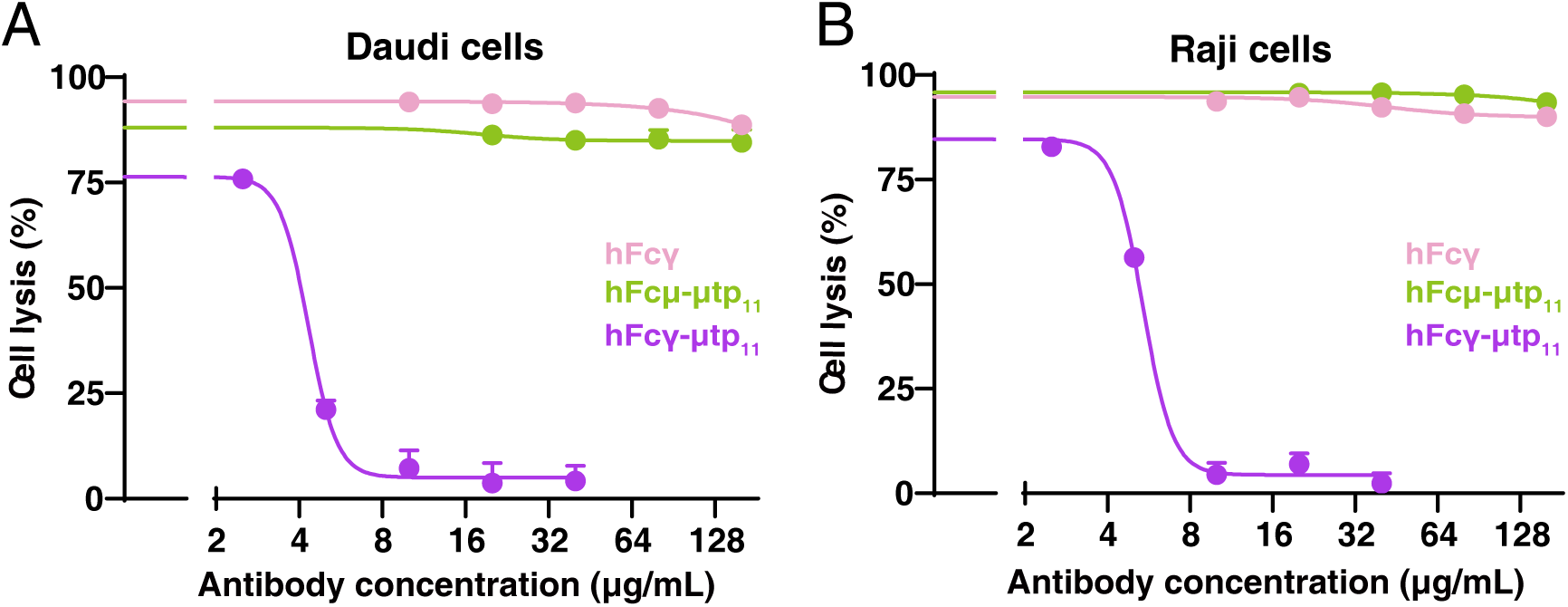
Engineered IgG-Fc hexamer with μtp_11_ blocks RTX-IgG mediated complement activation. (A) The IgG-Fc hexamer with μtp_11_ blocks RTX-IgG mediated complement activation in the Daudi cells. The IgM-Fc hexamers with μtp_11_ and IgG-Fc monomers exhibit no inhibitory effect. Data were plotted as the mean ± 1 s.d. *n* = 3 biological replicates. Source data are provided in the Source Data file. (B) The IgG-Fc hexamer with μtp_11_ blocks RTX-IgG mediated complement activation in the Raji cells. Data were plotted as the mean ± 1 s.d. *n* = 3 biological replicates.

## Discussion

Here we show that Xenopus IgX can form hexamers independently of the J-chain. We further demonstrated that by engineering the 11-amino acid IgX tailpiece onto human IgM, or by truncating the natural 18-amino acid tailpiece of IgM to 11-16 residues, the formation of IgM hexamers can be promoted. From a mechanistic perspective, having fewer than 11 residues in the tailpiece may compromise the integrity of the tailpiece β-strand, potentially destabilizing the IgM hexamer. Conversely, a tailpiece longer than 16 residues may hinder hexamer formation due to steric hindrance. Specifically, the penultimate Cys575 participates in disulfide bond formation between adjacent Fcμ units, while the terminal Tyr576 has a bulky side chain. Together, these factors could pose a challenge for accommodating twelve intact tailpieces within the confined central cavity of an Fcμ hexamer.

IgM antibodies have demonstrated potential for therapeutic applications(*48*). Naturally, IgM antibodies typically exhibit low affinity and specificity; however, grafting variable domains from affinity-matured IgG onto the IgM scaffold significantly enhances binding avidity and potency. For instance, converting a SARS-CoV-2 neutralizing IgG into an IgM format has markedly increased antibody potency and breadth(*49*). This engineered IgM can also be administered intranasally, making it a promising candidate for the prevention and treatment of SARS-CoV-2 and other respiratory pathogens. Other biologics utilizing the IgM scaffold have been developed, including an IgM-like ACE2 that broadly neutralizes SARS-CoV-2 variants and can be administered via aerosol inhalation(*50*), as well as an IgM-based multivalent nanobody platform called Adaptive Multi-Epitope Targeting with Enhanced Avidity (AMETA)(*51*). Our study offers straightforward strategies for engineering IgM hexamers, using either a shortened version of IgM’s own tailpiece or the tailpiece from IgX, which could enhance the efficacy and utility of these IgM-dependent biologics. It should be noted that despite their identical lengths, μtp_11_ and the 11-amino acid χtp differ in their sequences. In particular, μtp_11_ contains a more rigid Pro559 instead of a Ser in χtp, and an Asn563-Val564-Ser565 motif that leads to glycosylation at Asn563, unlike the Asn-Val-Asn sequence in χtp that does not support this modification (Fig. 2A). These differences do not seem to affect the ability of μtp_11_ to facilitate IgM hexamer formation; however, the impact of χtp and μtp_11_ on the efficiency of hexamer production awaits further investigation.

Importantly, by fusing a shortened IgM tailpiece onto IgG, we have also enabled IgG to assemble into a hexamer. The IgG hexamers exhibit potent complement-activating activities, potentially enhancing their application for therapeutic purposes(31). Compared to strategies involving the fusion of the 18-amino acid μtp tailpiece(32, 33), or the C575S and V567I/A572G variants with the same length(34, 36), our strategy could facilitate more efficient hexamer formation. As a proof of concept, our engineered RTX-Fcγ-μtp_11_ exhibited robust complement-activating properties, while the Fcγ-μtp_11_ fragments effectively blocked the CDC activity of the RTX IgG in a competitive manner. Collectively, these findings establish a foundation for an alternative strategy to generate IgG or IgG-Fc hexamers for therapeutic applications. Certainly, the half-life of our engineered IgG and Fcγ with μtp_11_ are likely shorter than that of wild-type counterparts, due to hindered interaction with FcRn(*52*). Similar challenges would also arise in other engineering strategies that that result in the formation of stable IgG hexamers, such as CSL777 (35) and Fcγ_Innovent_ (36). On the other hand, a shorter half-life may be advantageous in situations where toxicity is a concern. Apparently, significant further development is necessary to enhance usability of these molecules, particularly in relevant contexts. In contrast, we anticipate that the half-life of our engineered IgM would be comparable to that of wild-type IgM, as there is no FcRn-like receptor for IgM.

## Materials and Methods

### Cell culture

HEK293F cells (Thermo Fisher, 11625019) were cultured using SMM 293-TI medium (Sino Biological) in a humidified shaker at 37 °C with 5% CO_2_ and 55% humidity. OCI-Ly10 cells (RRID: CVCL_8795), Raji cells, and Daudi cells were originally purchased from the American Type Culture Collection and cultured in RPMI-1640 medium (Gibco) supplemented with 10% FBS (PAN Seratech) and 1% penicillin-streptomycin (Gibco) in a humidified incubator at 37 °C with 5% CO_2_.

### Protein expression and purification

Codon-optimized DNA fragments encoding the Fcχ (residues 240-584, UniProtKB: Q6INK3) or Fcχ^cχ3-cχ4-χtp^ (residues 360-584) were cloned into the modified pcDNA vector with an N-terminal IL-2 signal peptide followed by an 8×His tag. The constructs were transiently transfected into HEK293F cells using polyethylenimine (Polysciences) and cultured for 4 days. The protein was retrieved from the conditioned medium using the Ni-NTA affinity resin (Smart Lifesciences) and then further purified using a Superose 6 Increase column (GE Healthcare) in the final buffer containing 20mM HEPES, pH 7.2, and 150 mM NaCl.

For Fcμ with various tailpiece segments, the DNA fragments encoding Fcμ (residues 229-576, UniProtKB: P0DOX6) and its variants with χtp or different lengths of μtp were cloned into a modified pcDNA vector with a N-terminal IL-2 signal peptide followed by a twin-strep tag. The proteins were expressed similarly and purified first using the Strep-Tactin resin (Smart Lifesciences) and then further purified using a Superose 6 Increase column.

To generate Fcγ-μtp_11_ chimera protein, the truncated μtp_11_ segment was introduced into the C-terminus of Fcγ (residues 218-449, UniProtKB: P0DOX5) and the C-terminal region of Fcγ (SLSPGK) was replaced with the corresponding region in Fcμ (DKSTGK). The DNA fragment encoding Fcγ-μtp_11_ was cloned into a modified pcDNA vector with an N-terminal IL-2 signal peptide followed by a Flag tag. The construct expressing Fcγ-μtp_11_ was transfected into HEK293F cells. After 4 days, the conditioned medium was incubated with the anti-Flag M2 affinity gel (A2220, Sigma-Aldrich) and eluted using the binding buffer supplemented with 200 μg/mL 3× Flag peptide (NJP50002, NJPeptide). The protein was then purified using a Superose 6 Increase column in buffer containing 20 mM HEPES, pH 7.2, and 150 mM NaCl. Purified proteins were examined by reduced SDS-PAGE or native-PAGE and Coomassie staining.

### Sedimentation velocity analytical ultracentrifugation (SV-AUC)

The SV-AUC experiments were conducted using a 12-mm charcoal-filled Epon centerpiece (Beckman, 392778) and a four-hole An60 Ti rotor spinning at 50,000 rpm in the Optima AUC analytical-ultracentrifuge (Beckman Coulter) with absorbance detection at 280 nm at 16 °C. The buffer solution consisted of 25 mM Tris-HCl, pH 7.4, and 150 mM NaCl. The data were analyzed using SEDFIT software to determine the sedimentation coefficient distribution *c*(*s*).

### Cryo-EM sample preparation and data collection

The protein samples were concentrated to 1.2-1.5 mg/mL and then treated with 0.05% glutaraldehyde (Sigma) at 20 °C for 10 min. 4 μL aliquots of the crosslinked samples were applied onto glow-discharged holey carbon gold grids (Quantifoil, R1.2/1.3) using a Vitrobot Mark IV at 4 °C with 100% humidity. The blotting time was 1.0 s, followed by a waiting time of 10 s. The grids were then plunged into liquid ethane. The grids were screened using a 200 kV Talos Arctica microscope equipped with a Ceta camera. Data collections were performed using a 300 kV Titan Krios G3 electron microscope with a K3 Summit direct detection camera, or a 300 kV Titan Krios G4 electron microscope with a Falcon 4 camera. Data were collected using the EPU software (E Pluribus Unum, Thermo Fisher) and the defocus ranges were set from −1.0 to - 1.4 μm.

### Cryo-EM data processing and model building

Raw movie frames were motion-corrected using MotionCor2 (v1.4.4)(*53*). The contrast transfer function (CTF) parameters were estimated using Gctf (v1.06)(*54*). Subsequent data processing was carried out using cryoSPARC (v3.2)(*55*). Summed images were screened manually, and particles were picked using the blob picker. Templates were generated by 2D classification, and the template-picking particles were subjected to several rounds of 2D classification to exclude inaccurate particles and then further subjected to ab-initio reconstruction and heterogeneous refinement. The particles retained from heterogeneous refinement were subjected to homogeneous and non-uniform refinement to generate the final 3D reconstruction.

To improve the density of the tp region, the Fcχ^Cχ3–Cχ4–χtp^ and Fcμ-μtp_11_ particles were imported into RELION (v5.0), where C6 symmetry 3D refinement was performed, followed by C6 symmetry expansion to generate all symmetry-related views. Subsequently, 3D classification without any alignment was performed using a local mask focused on the tp region.

The local resolution map was analyzed using ResMap(*56*) and displayed using UCSF ChimeraX(*57*). The structure models were then adjusted using Coot(*58*) and refined using the real-space refinement in Phenix(*59*).

### Complement-dependent cytotoxicity assay

To generate engineered anti-CD20 IgM and IgG variants, the heavy chain DNAs of the antigen-binding fragments of rituximab (RTX) were mounted upstream of Fcμ, Fcμ-μtp_11_, and Fcγ-μtp_11_ in the pcDNA vector. The resulting plasmids were transfected into HEK293F cells together with the corresponding light chain, with or without J-chain expression plasmids using a 1:1:2 or 1:1 ratio. The proteins were then isolated from the conditioned medium using the Ni-NTA or the anti-Flag M2 affinity gel and then the Superose 6 Increase column as described above. All proteins were analyzed by reduced SDS-PAGE or native-PAGE and Coomassie staining.

The complement-dependent cytotoxicity assay was conducted using various B lymphoma cell lines, including OCI-Ly10, Daudi, and Raji. The OCI-Ly10 cells were cultured with standard FBS and used directly in the assay; whereas the Daudi and Raji cells were cultured for several days in RPMI-1640 medium supplemented with 10% heat-inactivated FBS prior to the assay. The RTX-Fcμ or RTX-Fcγ proteins were incubated with equal volumes of cell cultures (∼20,000 cells) and 12% normal human serum complement (Quidel) sequentially, and then transferred into a 96-microwell plate. After 8 h of incubation at 37 °C, 50 μl of CellTiter-Glo reagent (Promega, G7572) was added to each well and incubated for 10 min at room temperature. Luminescence was measured using a Cytation 5 cell imaging multimode reader (BioTek). The data were analyzed by plotting the luminescence units against concentrations of the proteins in GraphPad Prism using a 4-parameter curve-fit.

To verify the inhibitory effect of engineered Fcγ-μtp_11_ hexamers, Fcμ-μtp_11_ hexamers, and Fcγ monomers for RTX-Fcγ-mediated classical complement activation, RTX-Fcγ proteins (1.2 μg/mL) were mixed with serially diluted Fcγ or Fcμ proteins. The resulting samples were further incubated with equal volumes of B lymphoma cell cultures (∼20,000 cells) and 12% normal human serum complement sequentially and then transferred into a 96-microwell plate. After 8 h of incubation at 37 °C, the samples were measured and analyzed as described as above.

## Acknowledgments

We are grateful to the Cryo-EM and the high-performance computing Platforms of Peking University and Changping Laboratory for their support with data collection and computation.

## Funding

This work received support from the National Natural Science Foundation of China (32325018) and the Qidong-SLS Innovation Fund to J.X. C.J. is supported by the Boya Postdoctoral Fellowship from Peking University.

## Author contributions

R.Z. and C.J. conducted protein design, purification, structural studies, and functional assays. S.L. assisted with protein expression and purification. R.Z. and C.J. coordinated the mouse analyses. J.X. conceived and supervised the project, and wrote the manuscript with contributions from all authors.

## Competing interests

R.Z., C.J., and J.X. are listed as inventors on a patent application that involves the development of IgM and IgG hexamers.

## Data and materials availability

The cryo-EM map and atomic coordinates of Fcχ^Cχ3–Cχ4–χtp^ hexamer, hFcμ-χtp hexamer, hFcμ-μtp_11_ hexamer, hFcγ-μtp_11_ hexamer have been deposited in the EMDB and PDB with accession codes EMD-64375, EMD-64374 (local map), and 9UO6; EMD-64372 and 9UO4; EMD-64371 and 9UO3; EMD-64373 and 9UO5; respectively.

---

**Fig. S1.**
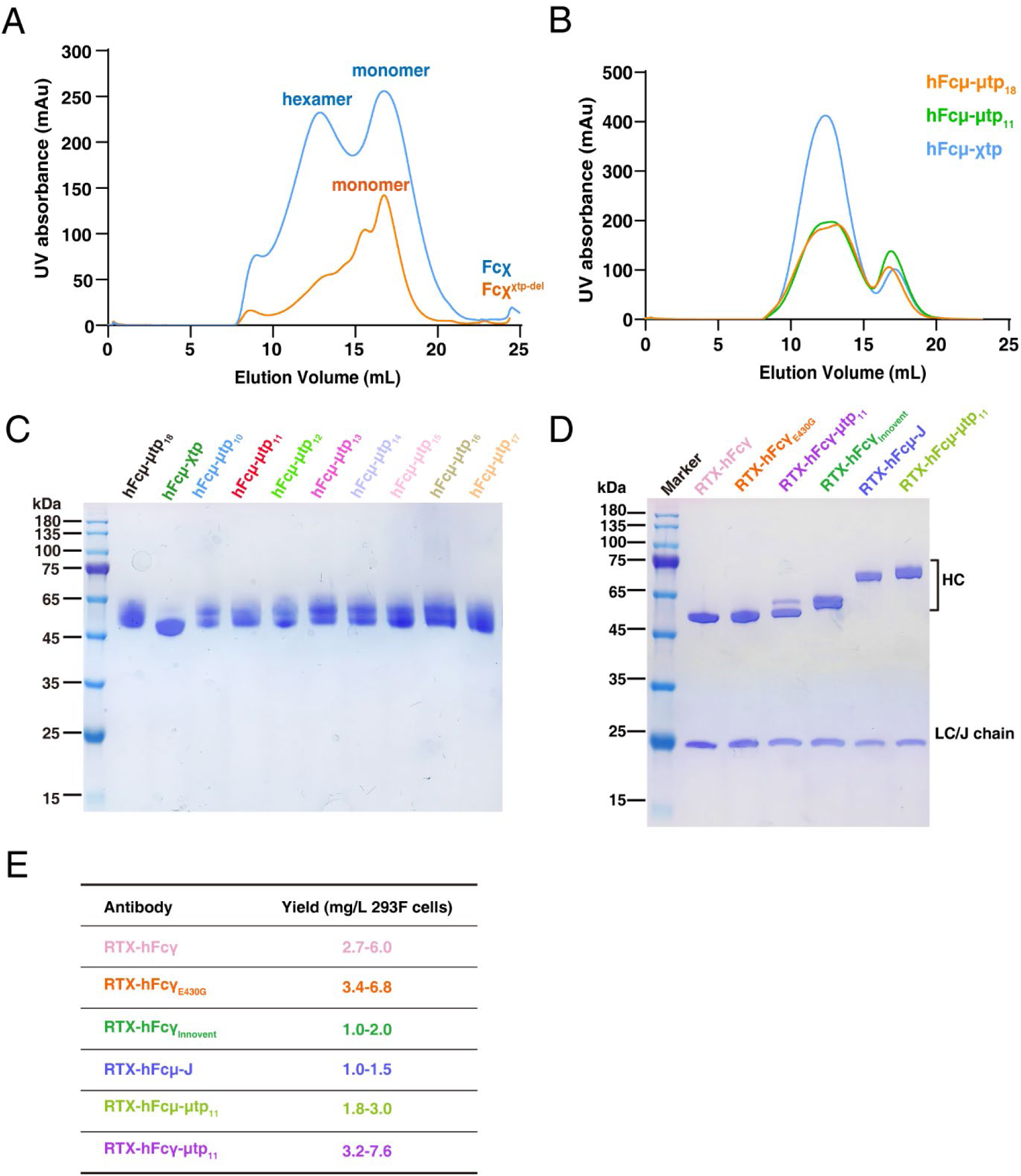
Protein purifications. A. Size exclusion chromatography (SEC) profiles of purified Fcχ and Fcχ_Δχtp_. Fcχ is shown in blue, and Fcχ_Δχtp_ is shown in orange. B. SEC profiles of purified Fcμ-μtp variants. C. SDS-PAGE analyses of purified Fcμ-μtp variants. D. SDS-PAGE analyses of the engineered RTX-Fcγ and RTX-Fcμ antibodies after purification. E. Purification yield (mg/L) of various Igs from HEK293F cells under our lab conditions.

**Fig. S2.**
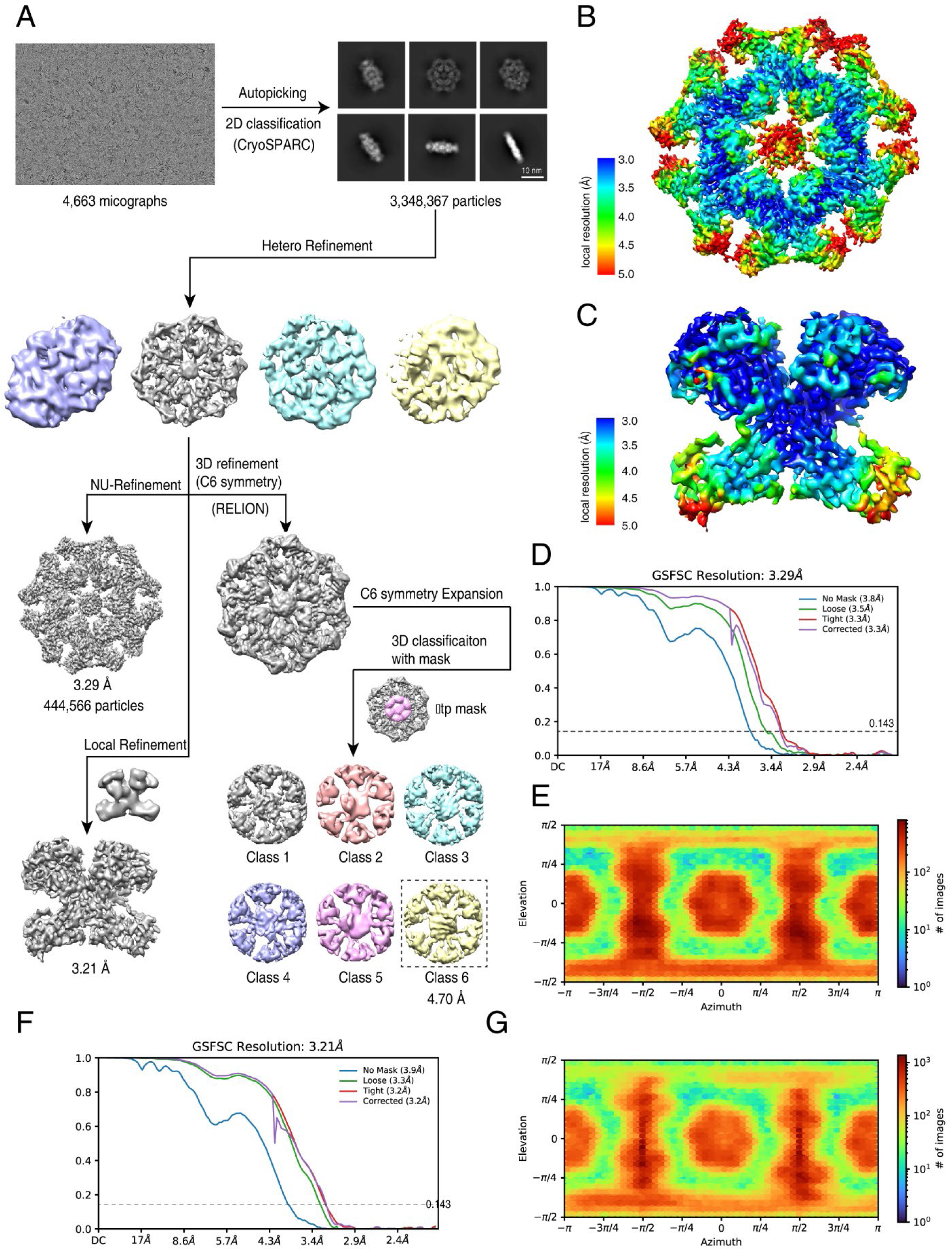
Cryo-EM 3D reconstruction of the Fcχ^Cχ3–Cχ4–χtp^ hexamer. A. Flowchart of cryo-EM data processing of Fcχ^Cχ3-Cχ4-χtp^. B. Resolution estimations of the overall map of Fcχ^Cχ3-Cχ4-χtp^. C. Resolution estimations of the local map around the Fcχ-Fcχ interface of Fcχ^Cχ3-Cχ4-χtp^. D. FSC curves with estimated resolutions of Fcχ^Cχ3-Cχ4-χtp^. E. Angular distribution of the Fcχ^Cχ3-Cχ4-χtp^ particles used in the final 3D reconstruction. F. FSC curves with estimated resolutions for the local map. G. Angular distribution of the Fcχ^Cχ3-Cχ4-χtp^ particles used in final 3D reconstruction for the local map.

**Fig. S3.**
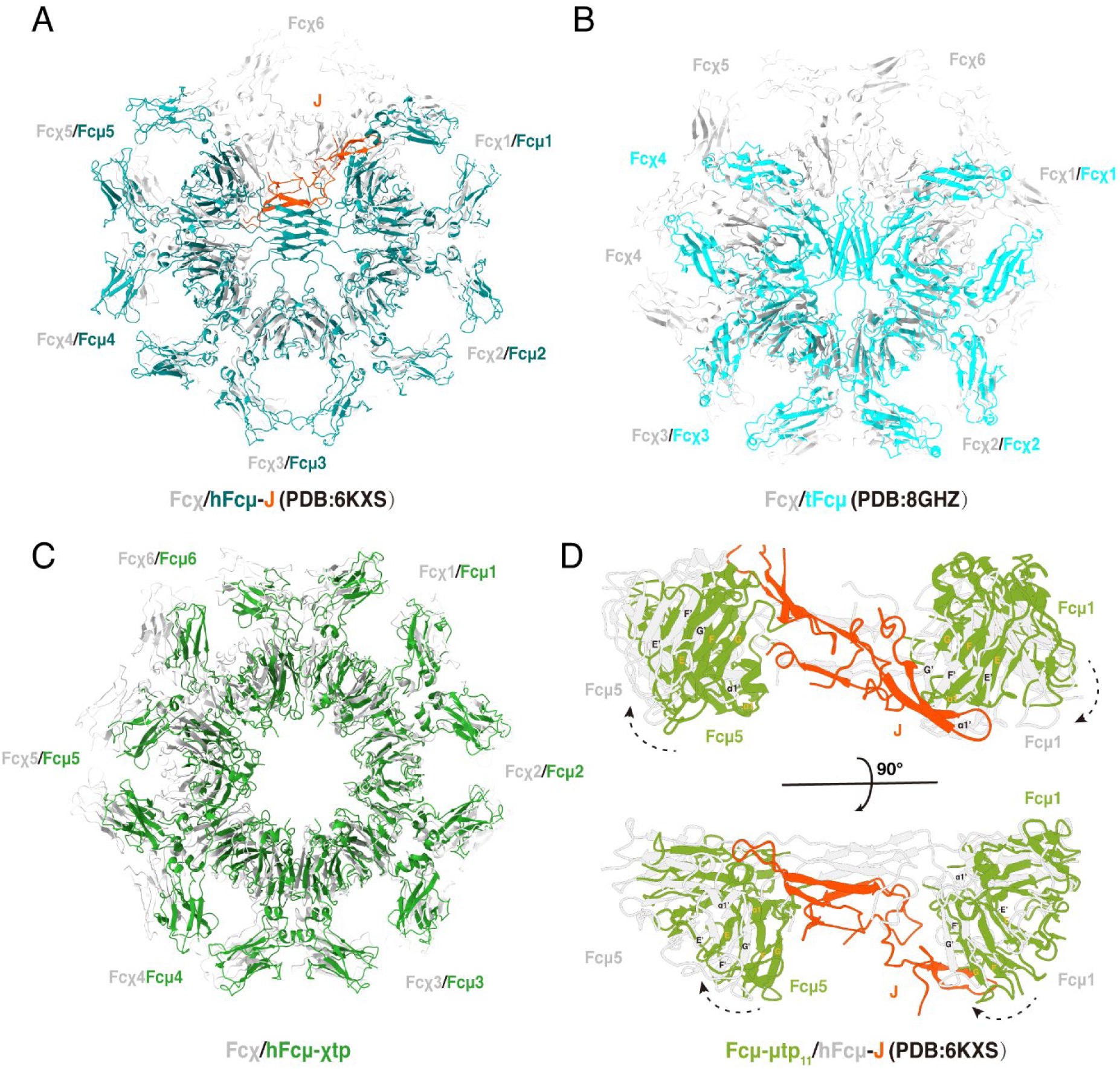
Structural comparisons. A. Structural overlay of the Fcχ hexamer and the Fcμ-J pentamer. Fcχ is shown in white, whereas Fcμ is shown in teal. J-chain in Fcμ-J is shown in orange. B. Structural overlay of the Fcχ hexamer and the teleost Fcμ tetramer. Fcχ is shown in white, whereas tFcμ is shown in cyan. C. Structural comparison of the Fcχ hexamer and the Fcμ-χtp hexamer. Fcχ is shown in white, and Fcμ is shown in green. D. Structural comparison of Fcμ-μtp_11_ hexamer and the Fcμ-J pentamer at the Fcμ1-Cμ4 and Fcμ5-Cμ4 regions. The Fcμ molecules in the Fcμ-μtp_11_ hexamer and the Fcμ-J pentamer are shown in green and white, respectively. J-chain is colored in orange.

**Fig. S4.**
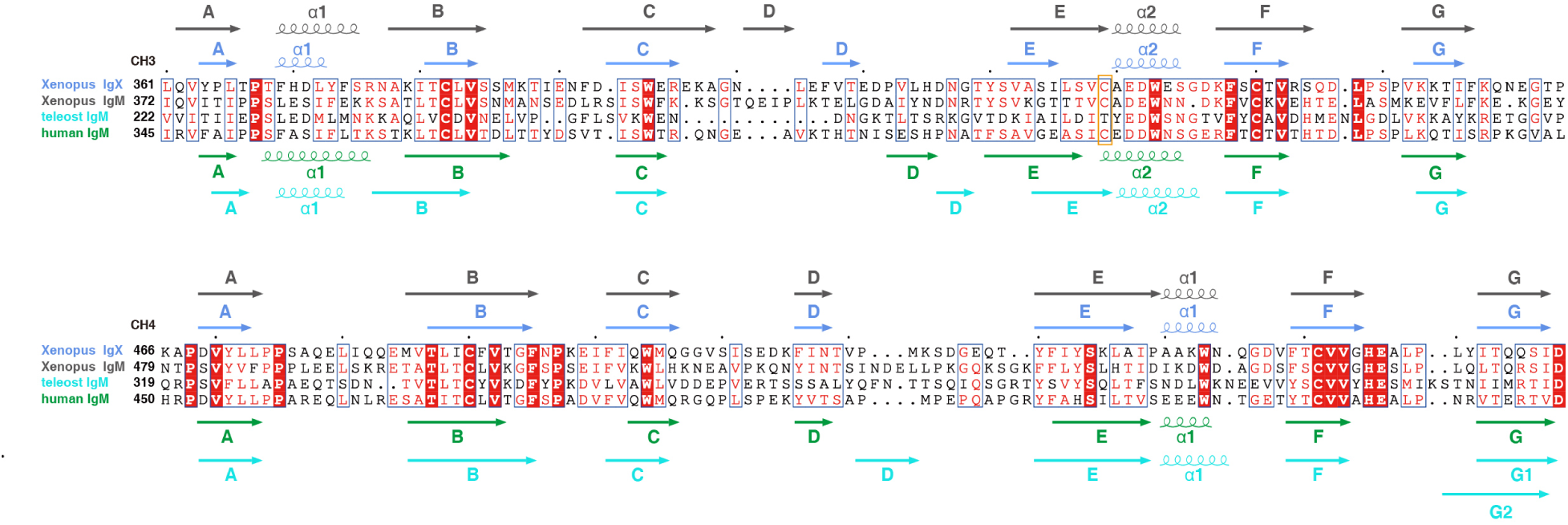
Sequence alignments of the CH3-CH4 regions of Xenopus IgX, Xenopus IgM, teleost IgM, and human IgM. Secondary structures are depicted above and below the sequence blocks, color-coded to match the labels of the respective molecules. The orange rectangle highlights Cys430 in the Cχ3 domain of IgX, along with the corresponding Cys residues in Xenopus and human IgM. In contrast, a comparable Cys residue is notably absent in the teleost IgM.

**Fig. S5.**
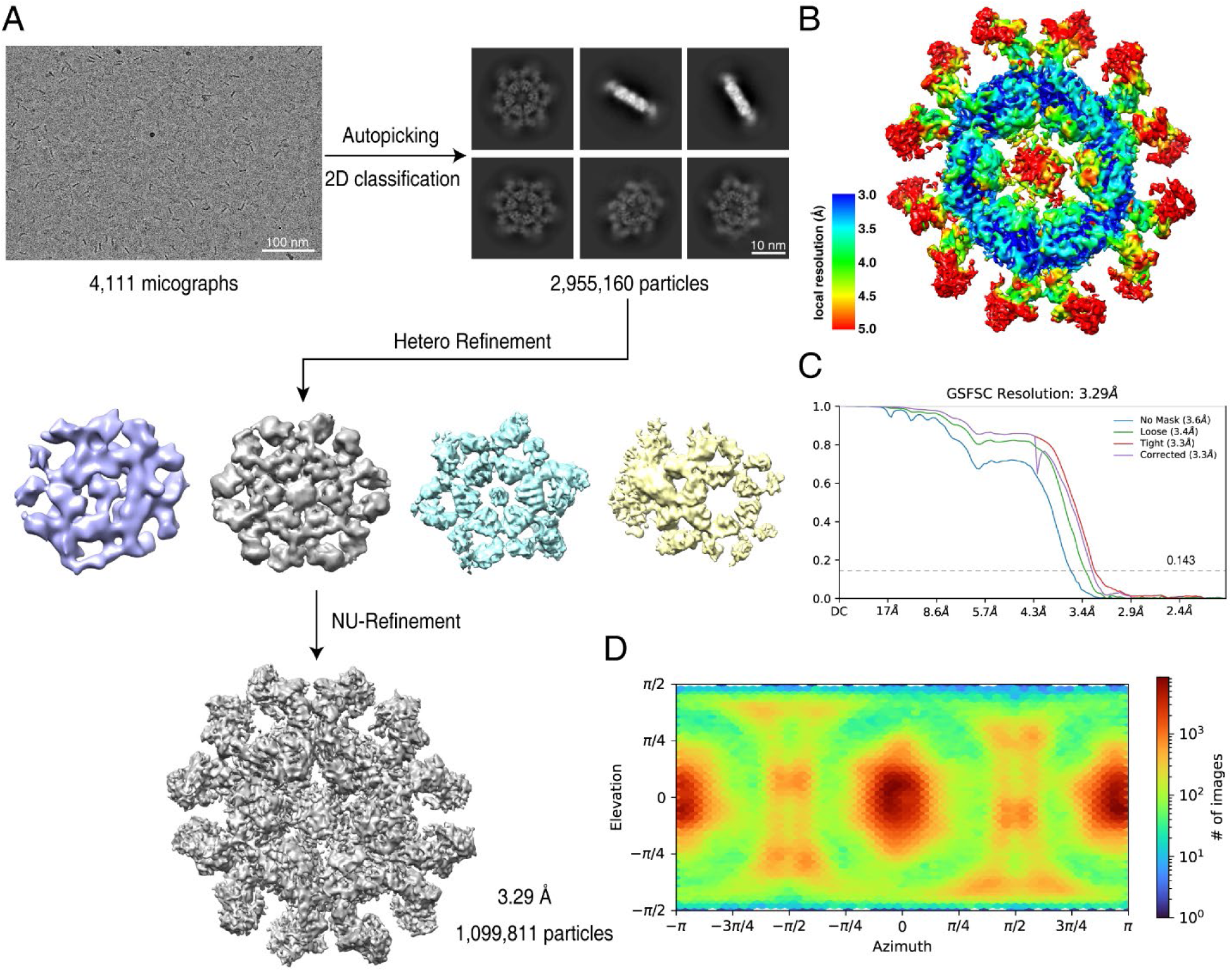
Cryo-EM 3D reconstruction of the Fcμ-χtp chimera. A. Flowchart of cryo-EM data processing of Fcμ-χtp. B. Resolution estimations of the overall map of Fcμ-χtp. C. FSC curves with estimated resolutions of Fcμ-χtp. D. Angular distribution of the Fcμ-χtp particles used in final 3D reconstruction.

**Fig. S6.**
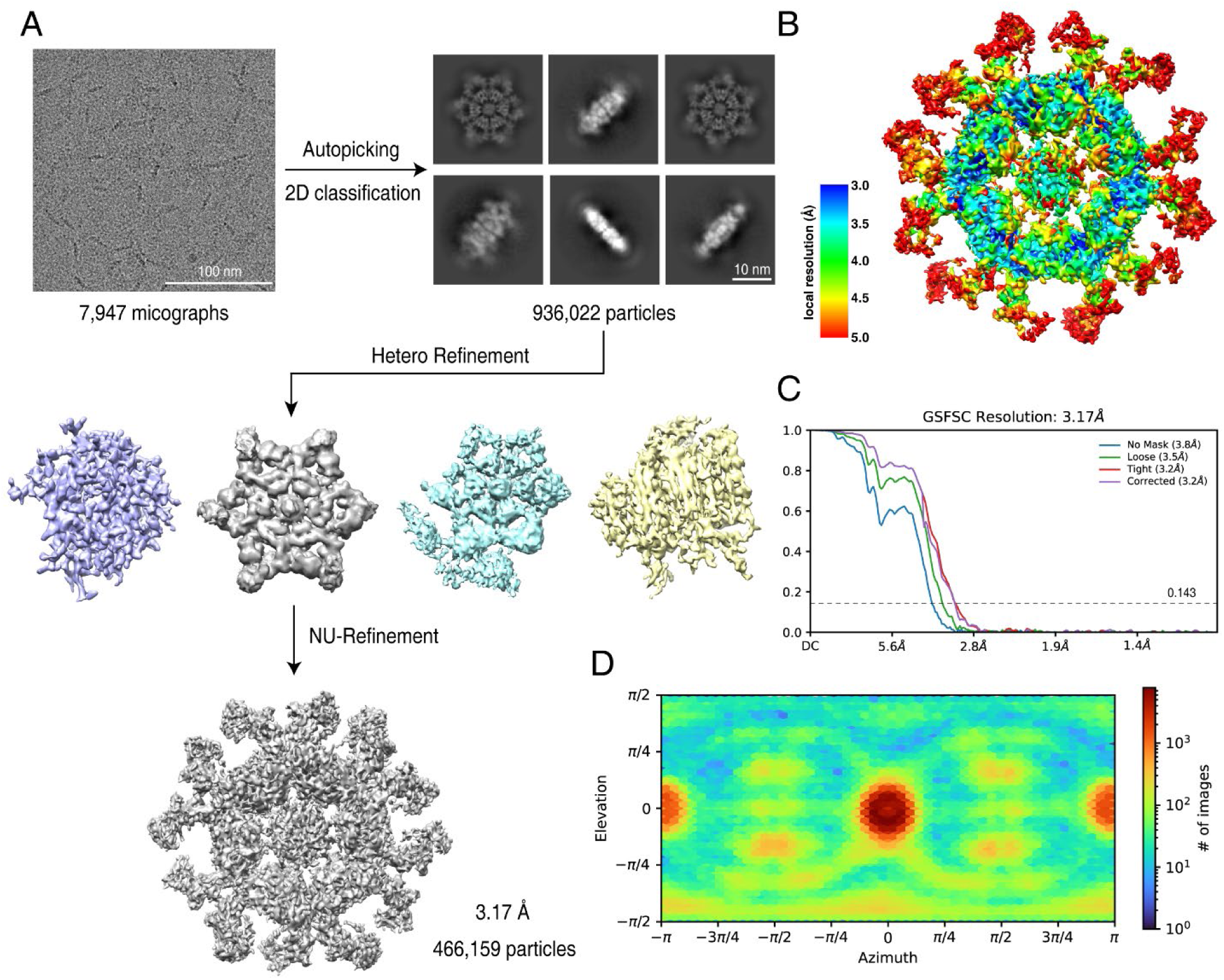
Cryo-EM 3D reconstruction of the Fcμ-μtp_11_ hexamer. A. Flowchart of cryo-EM data processing of Fcμ-μtp_11_. B. Resolution estimations of the overall map of Fcμ-μtp_11_. C. FSC curves with estimated resolutions of Fcμ-μtp_11_. D. Angular distribution of the Fcμ-μtp_11_ particles used in final 3D reconstruction.

**Fig. S7.**
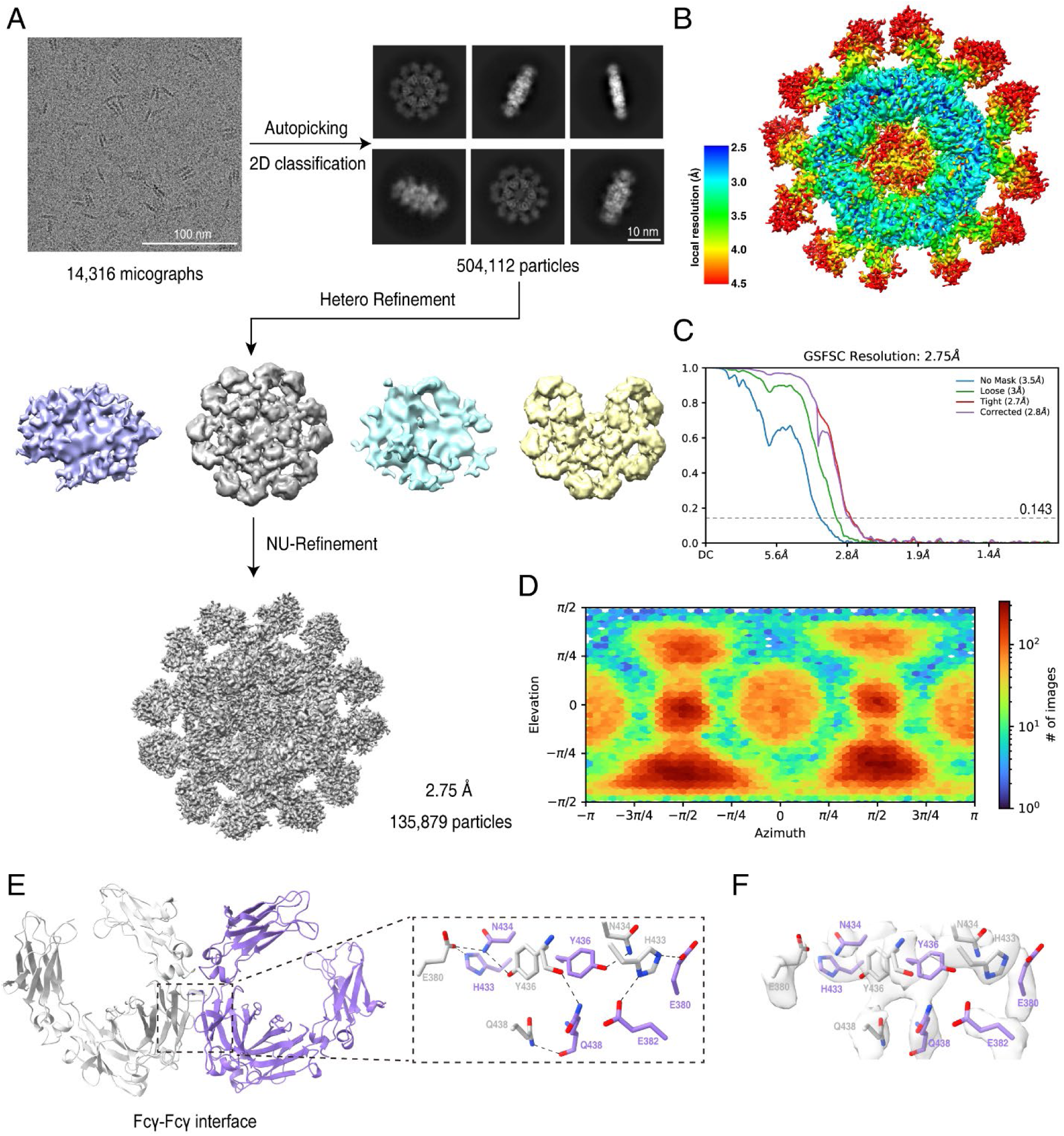
Cryo-EM 3D reconstruction of the Fcγ–μtp_11_ hexamer. A. Flowchart of cryo-EM data processing of Fcγ-μtp_11_. B. Resolution estimations of the overall map of Fcγ-μtp_11_. C. FSC curves with estimated resolutions of Fcγ-μtp_11_. D. Angular distribution of the Fcγ-μtp_11_ particles used in final 3D reconstruction. E. The interactions between two adjacent Fcγ domains. F. Density maps of the interactions at Fcγ-Fcγ interface.

**Table S1.** Cryo-EM data collection, refinement and validation statistics.

|  | **Fcχ**<br>(EMDB-64375,<br>EMDB-64374)<br>(PDB 9UO6) | **hFcμ-χtp**<br>(EMDB-64372)<br>(PDB 9UO4) | **hFcμ-μtp<sub>11</sub>**<br>(EMDB-64371)<br>(PDB 9UO3) | **hFcγ-μtp<sub>11</sub>**<br>(EMDB-64373)<br>(PDB 9UO5) |
| --- | --- | --- | --- | --- |
| **Data collection and processing** |  |  |  |  |
| Voltage (kV) | 300 |  | 300 |  |
| Microscope | Titan Krios G3 |  | Titan Krios G4 |  |
| Camera | K3 Summit |  | Falcon 4 |  |
| Magnification (calibrated) | 81,000× | | 215,000× | |
| Electron exposure (e<sup>-</sup>/Å<sup>2</sup>) | 61.04 | 60 | 60 |  |
| Defocus range (μm) | -1.0 to -1.4 |  | -1.0 to -1.4 |  |
| Pixel size (Å) | 1.07 |  | 0.56 |  |
| Micrographs used | 4,663 | 4,111 | 7,947 | 14,316 |
| Symmetry imposed | C1 | C1 | C1 | C1 |
| Initial particle images | 3,463,703 | 2,955,160 | 936,022 | 504,112 |
| Final particle images | 444,566 | 1,099,811 | 466,159 | 135,879 |
| Overall map resolution at 0.143 FSC of masked reconstruction (Å) | 3.29 | 3.29 | 3.17 | 2.75 |
| Local map resolution at 0.143 FSC of masked reconstruction (Å) | 3.21 | / | / | / |
| Overall map sharpening B factor (Å<sup>2</sup>) | -119.5 | -122.9 | -91.4 | -65.9 |
| Local map sharpening B factor (Å<sup>2</sup>) | -107.2 | / | / | / |
| **Refinement** |  |  |  |  |
| Map-model CC |  |  |  |  |
| CC_mask | 0.77 | 0.58 | 0.58 | 0.72 |
| CC_box | 0.73 | 0.59 | 0.59 | 0.65 |
| CC_peaks | 0.67 | 0.50 | 0.45 | 0.60 |
| CC_volume | 0.76 | 0.56 | 0.55 | 0.69 |
| Model composition |  |  |  |  |
| Non-hydrogen atoms | 20076 | 19,980 | 19,980 | 20,244 |
| Protein residues | 2538 | 2,568 | 2,568 | 2,540 |
| Ligands | 0 | 0 | 0 | 0 |
| Bond lengths (Å) | 0.003 | 0.003 | 0.005 | 0.003 |
| Bond angles (°) | 0.695 | 0.700 | 0.756 | 0.674 |
| B factors (Å<sup>2</sup>) |  |  |  |  |
| Protein | 91.34 | 80.06 | 44.52 | 12.2 |
| Ligands | / | / | / | / |
| Validation |  |  |  |  |
| MolProbity score | 1.80 | 1.95 | 1.94 | 1.76 |
| Clashscore | 7.16 | 9.22 | 9.22 | 8.55 |
| Poor rotamers (%) | 0.13 | 0.00 | 0.35 | 0.30 |
| Ramachandran plot |  |  |  |  |
| Favored (%) | 93.83 | 92.61 | 93.00 | 95.67 |
| Allowed (%) | 6.17 | 7.39 | 7.00 | 4.33 |
| Disallowed (%) | 0.00 | 0.00 | 0.00 | 0.00 |
| Cβ outliers (%) | 0.00 | 0.00 | 0.00 | 0.00 |
| CaBLAM outliers (%) | 3.53 | 3.10 | 5.00 | 2.85 |

